## Supplementary material for "Large-scale characterization of drug mechanism of action using proteome-wide thermal shift assays": Figure 2 - source data 3

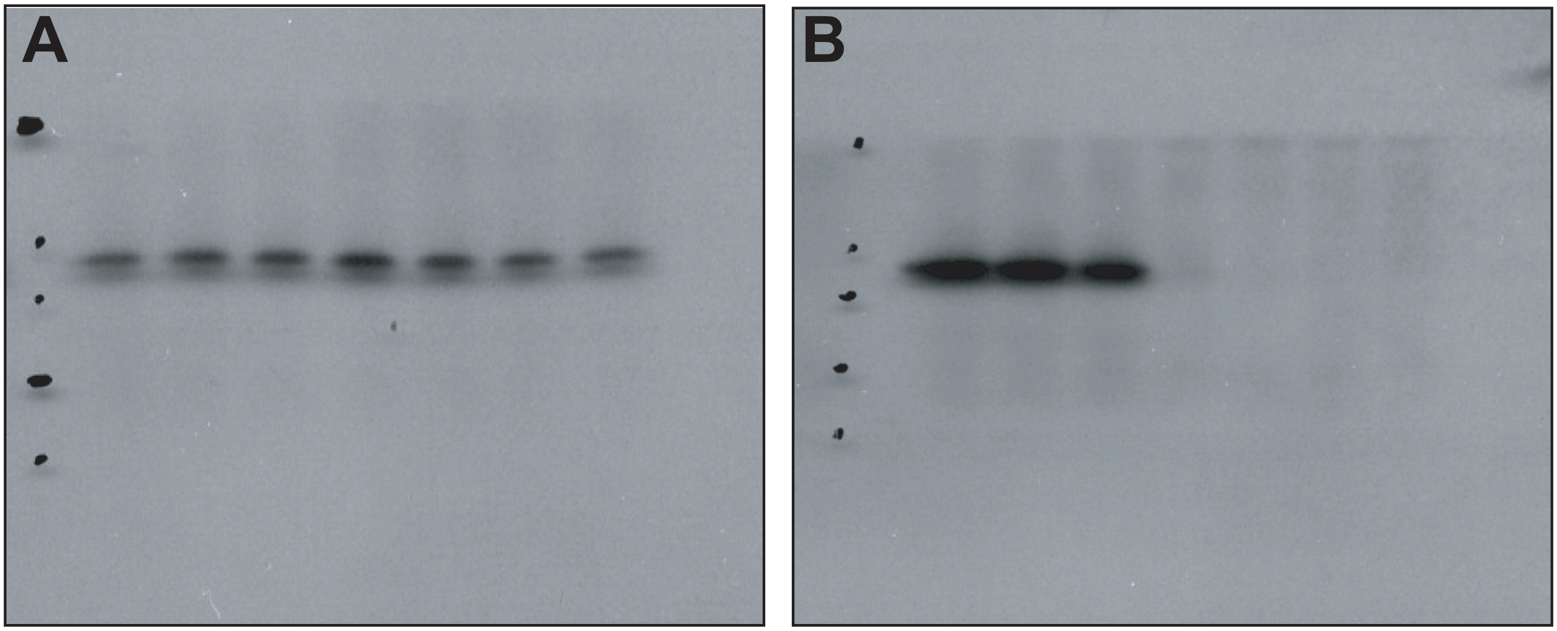


Figure 2F. Unedited scans of TCTP (A) and pTCTP (B).

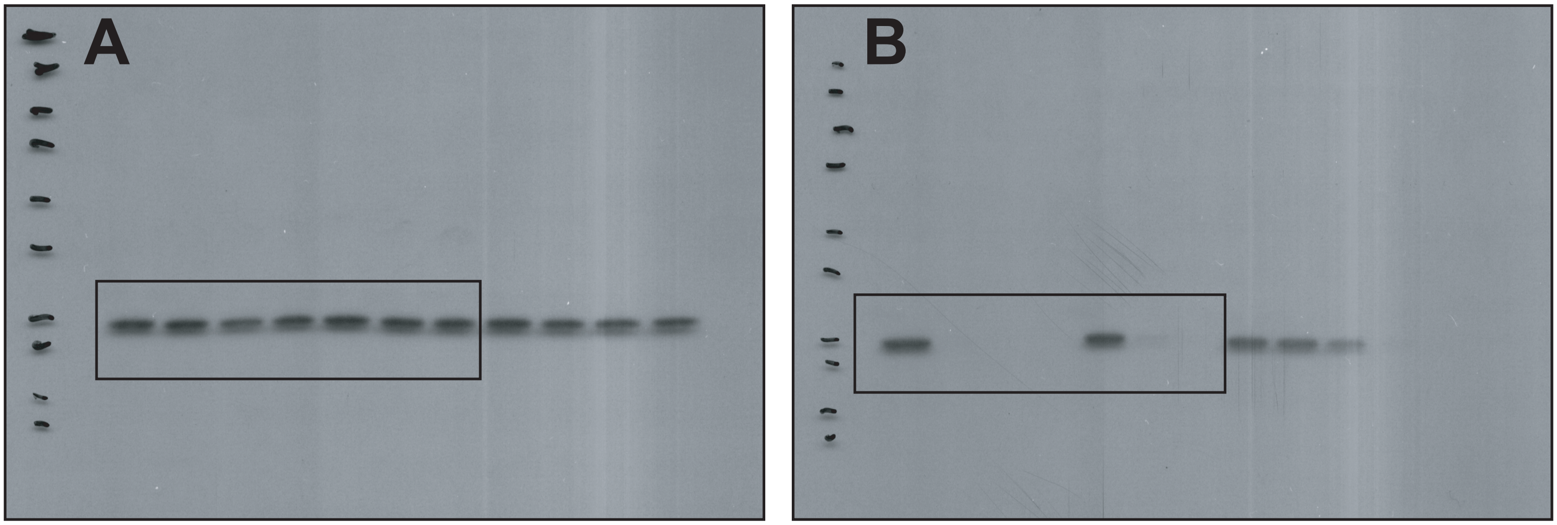


Figure 2 - figure supplement 1D. Unedited scans of TCTP (A) and pTCTP (B).

### Figure 2 - source data 3
