## Supplementary figures and images for "Large-scale characterization of drug mechanism of action using proteome-wide thermal shift assays"

### Figure 4 - source data 2

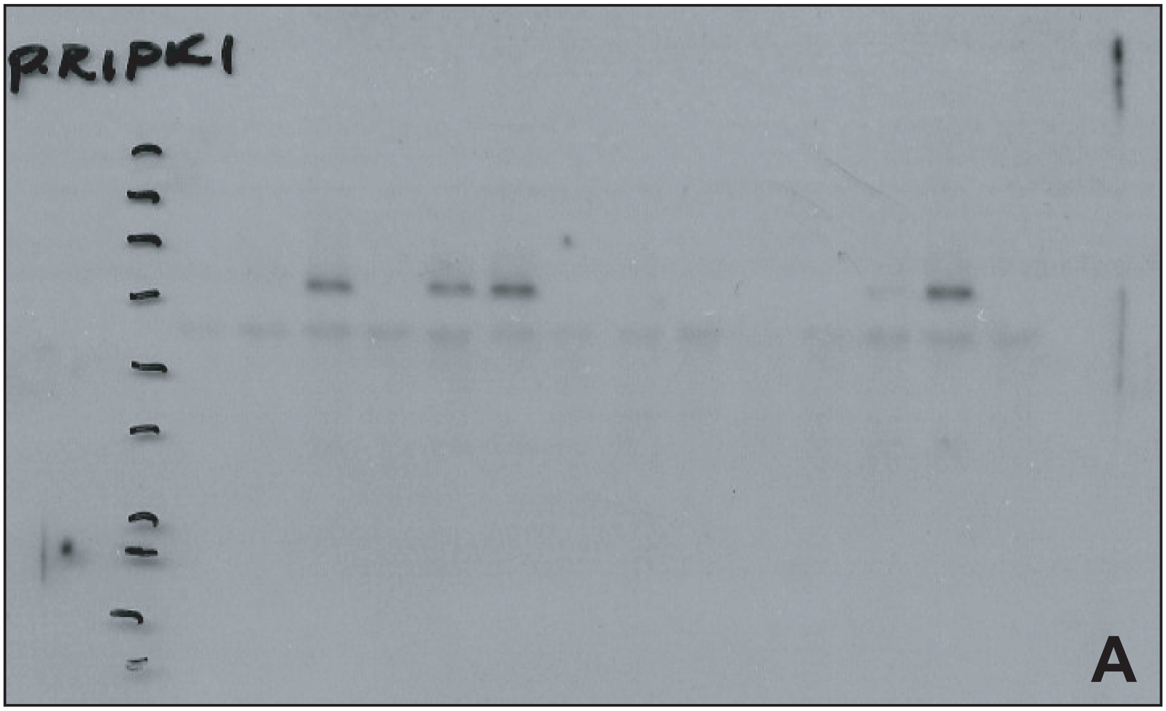


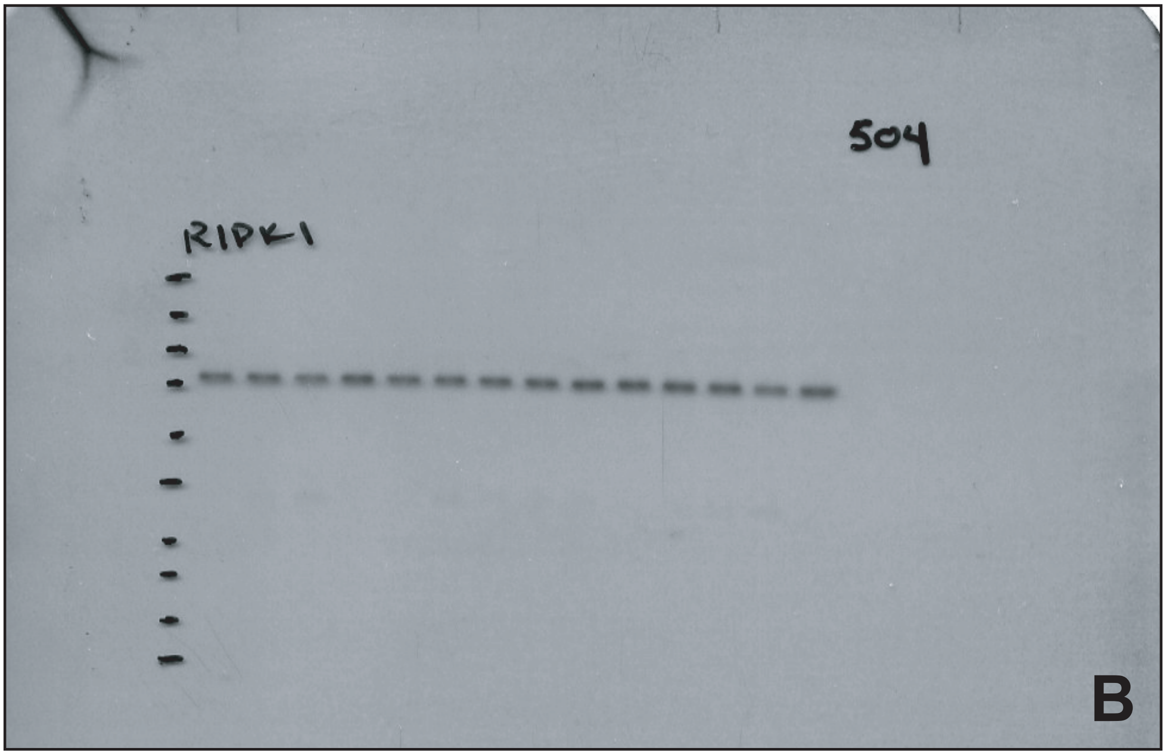


Figure 4G. Unedited scans of pRIPK1 (A) and RIPK1 (B).

#### Figure 4 - source data 2

### Figure 5 - source data 1

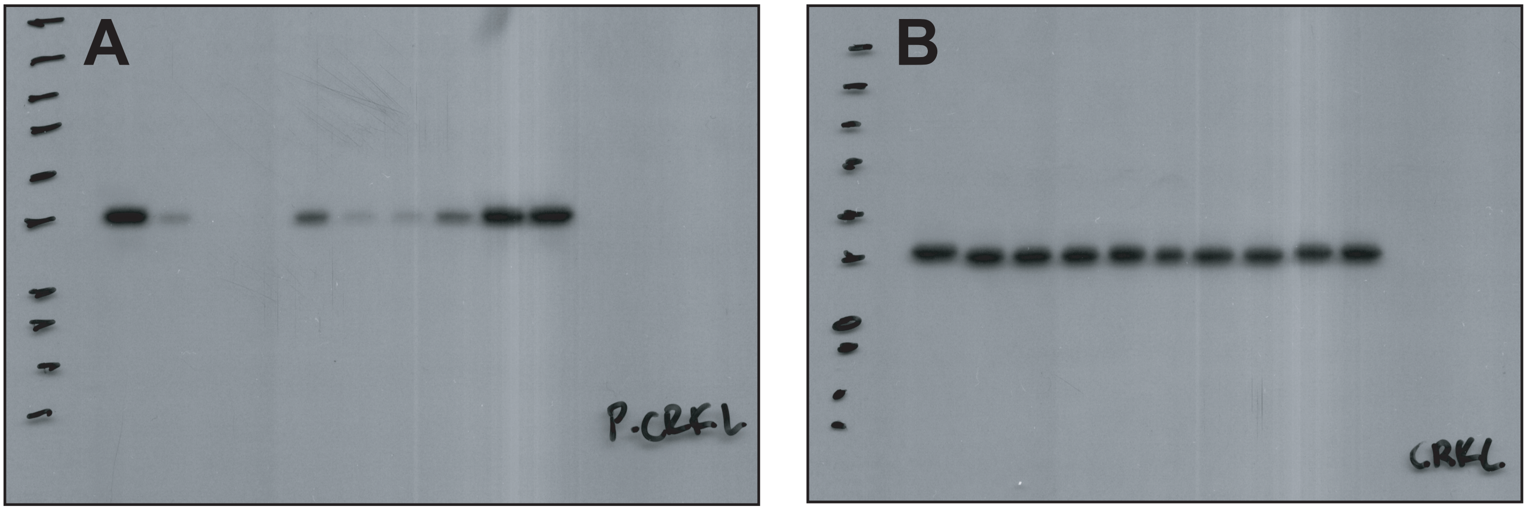


Figure 5C. Unedited scans of pCRKL (A) and CRKL (B).

#### Figure 5 - source data 1
